# Comparison of monomeric variants of StayGold

**DOI:** 10.1101/2024.02.28.582207

**Authors:** Satoshi Shimozono, Ryoko Ando, Mayu Sugiyama, Masahiko Hirano, Yusuke Niino, Atsushi Miyawaki

**Affiliations:** Laboratory for Cell Function Dynamics, RIKEN Center for Brain Science, 2-1 Hirosawa, Wako-city, Saitama 351-0198, Japan; Biotechnological Optics Research Team, RIKEN Center for Advanced Photonics, 2-1 Hirosawa, Wako-city, Saitama 351-0198, Japan; Department of Optical Biomedical Science, Institute for Life and Medical Sciences, Kyoto University. Kyoto 606-8507, Japan; Laboratory of Bioresponse Analysis, Institute for Life and Medical Sciences, Kyoto University. Kyoto 606-8507, Japan

## Abstract

StayGold is a bright and highly photostable fluorescent protein (FP) that forms an obligate dimer, thereby limiting its application as a soluble marker. On the basis of the structural information of this FP, we disrupted the dimerization to generate a monomeric variant, mStayGold, which inherits both the extremely high photostability and the high practical brightness of StayGold, for molecular fusion and membrane-targeting applications. Meanwhile, two other research groups have independently monomerized StayGold using different strategies. As a result, multiple StayGold monomers are currently available, creating confusion in the research community. In the present study, we investigated the three basic properties—photostability, brightness, and dispersibility—of these monomers by performing detailed side-by-side comparisons. This study highlights the difficulties of StayGold monomerization with emphasis on the tradeoff between photostability and brightness in FP technology.

## Introduction

The photobleaching of fluorescent proteins (FPs) is one of the most crucial problems in modern bioimaging. Through molecular cloning and mutagenesis studies on a wild-type FP from the jellyfish *Cytaeis uchidae*, we devised a solution. The engineered FP named StayGold is bright and hardly fades and can contribute to improving spatiotemporal resolution and extending the observation period^1^. As an obligate dimer, however, StayGold was initially used as a luminal soluble marker to visualize certain subcellular components, such as the endoplasmic reticulum (ER) and mitochondria.

In the Discussion section of our original paper^1^, we professed our actual engagement in directed evolution experiments that aimed to create mStayGold, a useful monomeric version of StayGold. In a subsequent study^2^, we determined the crystallographic structure of StayGold at 1.56 Å resolution. Similar to previous FP monomerization studies^3^, we identified mutational hotspots in the dimer interface on the basis of structural information (PDB ID, 8ILK). However, we soon realized that disrupting the dimerization while fully preserving the features of the original StayGold was difficult. Nevertheless, after multiple attempts with strict evaluation criteria for both photostability and brightness, we finally prepared excellent monomers, including the one that debuted very recently with the name mStayGold. To distinguish it from other variants, this monomer is herein tentatively termed “mStayGold(J),” where J stands for Japan (Fig. 1).

**Figure 1.**
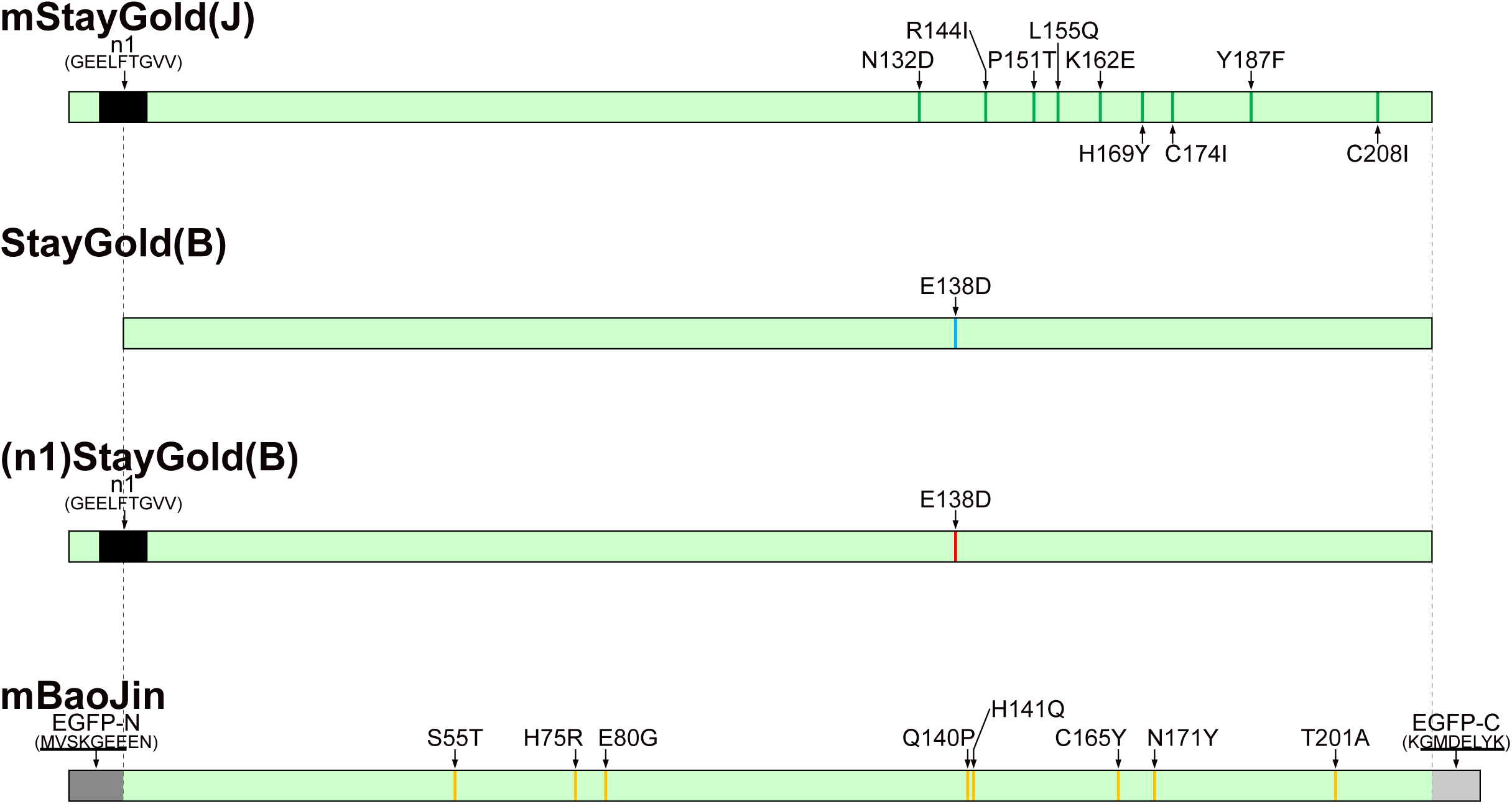
| StayGold monomers compared in the present study. Positions of mutations are indicated by vertical color bars. mStayGold(J), green; mStayGold(B), aqua; (n1)mStayGold, red; mBaoJin, orange. The n1 adaptor is shaded in black. The 7-residue N-terminus of EGFP (underlined) and 2 amino acids (EN) are shaded in dark gray. The 7-residue C-terminus of EGFP (underlined) and one amino acid (K) are shaded in light gray. The numbering is based on that of StayGold. See Supplementary Fig. 1.

On the other hand, Ivorra-Molla et al. determined the crystal structure of StayGold at 1.6 Å resolution (PDB ID, 8BXT)^4^. The British group investigated the monomericity of StayGold variants whose dimer interfaces had mutations predicted by the computer program PISA and reached a neat conclusion that a single mutation, E138D, can effectively monomerize StayGold to generate mStayGold (E138D) (Supplementary Note 1). By contrast, Piatkevich et al. performed elaborate genetic screening in bacteria to isolate StayGold monomers whose high brightness remained intact^5^. Numerous rounds of random mutagenesis gave rise to eight mutations in StayGold to generate another mStayGold, which now appears by the name of “mBaoJin” in the FPbase database.

Unfortunately, neither of the two research groups appreciated well our effort to extend the N– and C-termini of StayGold in the original study^1^. Both the head and tail of StayGold are relatively short, making this FP sensitive to C– and N-terminal fusions. Notably, the evolution of FPs toward useful tools has thus far involved substantial modifications of their N– and C-termini to develop fusion-tolerant FPs, such as the mFruit series^1,6^. Therefore, we strongly suggested the addition of n1 and c4 adaptors for C-terminal and N-terminal tagging, respectively, of the proteins of interest (POIs)^1,2^. In fact, mStayGold(J) has an n1 adaptor intrinsically and is recommended to be followed by a c4 adaptor for N-terminal tagging^2^. However, Ivorra-Molla et al. likely disregarded our suggestion and engineered StayGold by simply introducing a mutation into mStayGold (E138D)^4^ (Fig. 1). By contrast, Piatkevich et al. borrowed the N– and C-termini of EGFP (ref. 6) instead of using our adaptors for the extension of StayGold to make mBaoJin^5^ (Fig. 1).

To facilitate unbiased comparisons, we took utmost care to prepare constructs for analysis in the present study (Supplementary Fig. 1). First, we considered codon preference. Whereas the gene of mStayGold(J) has mammalian-preferred codons, the mStayGold (E138D) and mBaoJin genes have bacterial-preferred and jellyfish-original codons, respectively. Therefore, we generated a gene for mStayGold (E138D) with mammalian-preferred codons (Supplementary Fig. 2, top). The resultant variant is herein tentatively termed “mStayGold(B)” (Fig. 1). Also, we prepared a gene of mBaoJin with mammalian-preferred codons to maximize its expression level in mammalian cell types (Supplementary Fig. 2, bottom). Second, because the present study involved principally C-terminal tagging of POIs with StayGold monomers, we offered our n1 adaptor to mStayGold(B) to generate (n1)mStayGold(B) (Fig. 1). The use of these constructs enabled thorough side-by-side comparisons of different StayGold monomers developed independently by three research groups.

## Results

### Molecular brightness

We fused mStayGold(J), mStayGold(B), (n1)mStayGold(B), or mBaoJin to the C-terminus of a polyhistidine (His6) tag for expression in *Escherichia coli* (JM109 (DE3)) (Fig. 2a). Purified protein products were used for the determination of spectroscopic properties. mStayGold(J) absorbed light maximally at 499 nm, mStayGold(B) and (n1)mStayGold(B) at 496 nm, and mBaoJin at 497 nm (Fig. 2b). We determined the molar extinction coefficient (*ε*) by comparing the absorbance with the chromophore concentration (Fig. 2c). The *ε*s of mStayGold(J), mStayGold(B), (n1)mStayGold(B), and mBaoJin were calculated to be 163,900, 137,800, 141,100, and 135,700 M^-1^ cm^-1^, respectively (Table 1). The excitation and emission spectra of mStayGold(B), (n1)mStayGold(B), and mBaoJin were nearly the same. By comparison, the excitation and emission spectra of mStayGold(J) were red-shifted by 2–3 and 4–5 nm, respectively (Fig. 2d). This tiny but significant difference in the fluorescence spectra led us to characterize the spectral throughputs of the four StayGold monomers for correction of both excitation and emission collection efficiencies in the respective imaging systems (Supplementary Fig. 3). Lastly, we measured the absolute fluorescence quantum yield (QY_f_). mStayGold(J), mStayGold(B), (n1)mStayGold(B), and mBaoJin exhibited QY_f_ values of 0.83, 0.84, 0.85, and 0.80, respectively (Table 1). Regarding molecular brightness (the product of *ε* and QY_f_), mStayGold(J) exhibited the highest, followed by (n1)mStayGold(B), mStayGold(B), and mBaoJin.

**Figure 2.**
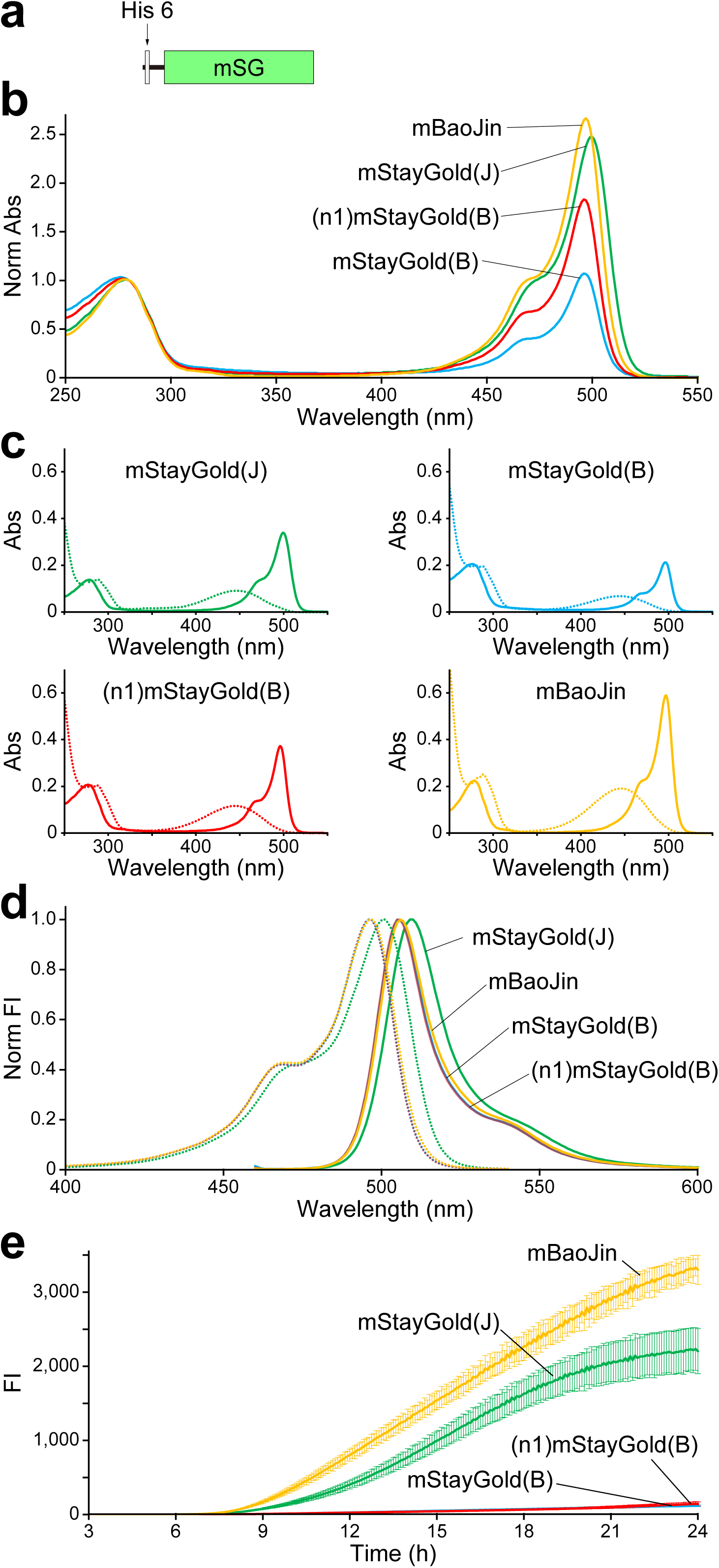
| Spectroscopic properties and bacterial expression brightness of StayGold monomers. **a**, Domain structure of His6-mSG. mSG: StayGold monomer. **b**, Absorption spectra of StayGold monomers, normalized against the peak at 280 nm. **c**, Absorption spectra of StayGold monomers before (solid line) and after (dotted line) denaturation with 0.1 M NaOH. The protein concentration was approximately 10 μM, and the light path length was 1 cm. **d**, Normalized excitation (dotted line) and emission (solid line) spectra of StayGold monomers. **e**, Fluorescence development in bacterial cells after plating at 37 L. The green fluorescence was first corrected for the respective spectral throughputs (Supplementary Fig. 3). Data points are shown as means ± s.d. (*n* = 6 different colonies from three independent experiments). **b–e**, Traces are indicated in green for mStayGold(J), aqua for mStayGold(B), red for (n1)mStayGold, and orange for mBaoJin. Abs: absorbance. FI: fluorescence intensity.

**Table 1.**
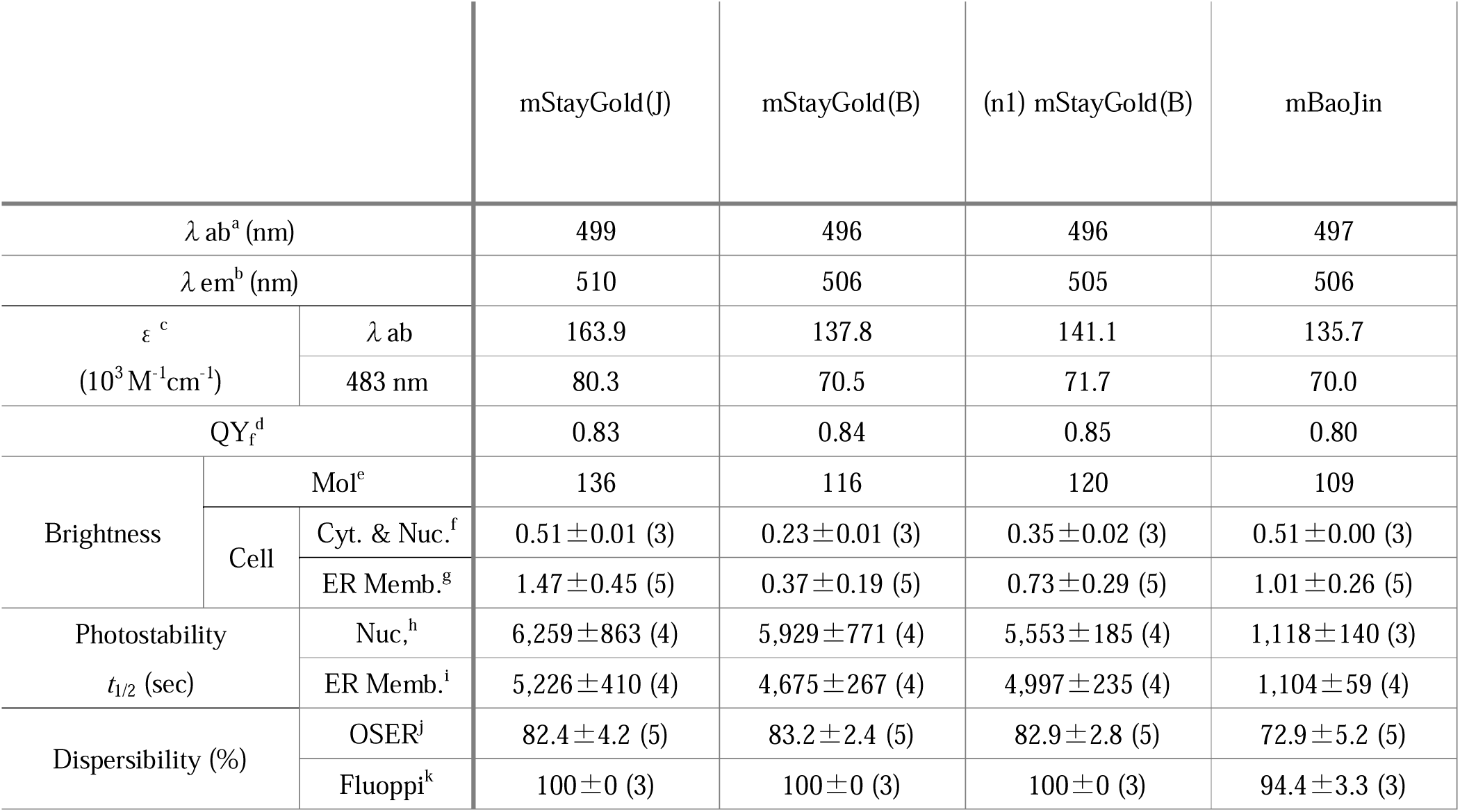
| Characteristics of StayGold monomers. ^a^ Absorbance maximum. ^b^ Emission maximum. ^c^ Absolute extinction coefficient *ε* at λab (top) and 483 nm (bottom). The measurement was based on the fact that after alkali denaturation of these FPs, the chromophore, containing a dehydrotyrosine residue conjugated to the imidazolone group, absorbs light maximally at 447 nm with a molar extinction coefficient of 44,000 M^−1^ cm^−1^. See Fig. 2c. *ε* (λab) values of mStayGold(B) and mBaoJin were reported to be 145 (ref. 4) and 124 (ref. 5), respectively. ^d^ Fluorescence quantum yield, QY_f_, measured using an absolute photoluminescence quantum yield spectrometer. QY_f_ values of mStayGold(B) and mBaoJin were reported to be 0.87 (ref. 4) and 0.81 (ref. 5), respectively. ^e^ Product of *ε* (λab) and QY_f_. This value reflects the molecular brightness of an FP. ^f^ Cellular brightness calculated from data shown in Fig. 3c. ^g^ Cellular brightness calculated from data shown in Fig. 5c. ^h^ Photostability calculated from data shown in Fig. 4. ^i^ Photostability calculated from data shown in Fig. 5e. ^j^ Percent whorl-free cells (Fig. 5d). ^k^ Percent punctum-free cells (Supplementary Fig. 6b). Values are means ± s.d. (*n* independent experiments). All values were measured by us in this study.

### Brightness in bacterial cells

We observed that, whereas bacterial colonies expressing mStayGold(J) or mBaoJin were very bright, those expressing mStayGold(B) or (n1)mStayGold(B) were dim. Fluorescence development after plating at 37 L was monitored simultaneously for 24 h (Fig. 2e). Apparently, mBaoJin matured faster than mStayGold(J). mStayGold(B) and (n1)mStayGold(B) were poorly expressed or remained immature during the observation period. We did not further characterize the brightness of these StayGold monomers in bacterial cells because their genes all had mammalian-preferred codons.

### Cellular brightness

We examined the actual brightness of the four StayGold monomers (mSGs) in living mammalian cells. We used a mammalian expression system (cotranslation via the T2A peptide) to correct the brightness of each monomer to that of mCherry (Fig. 3a).

**Figure 3.**
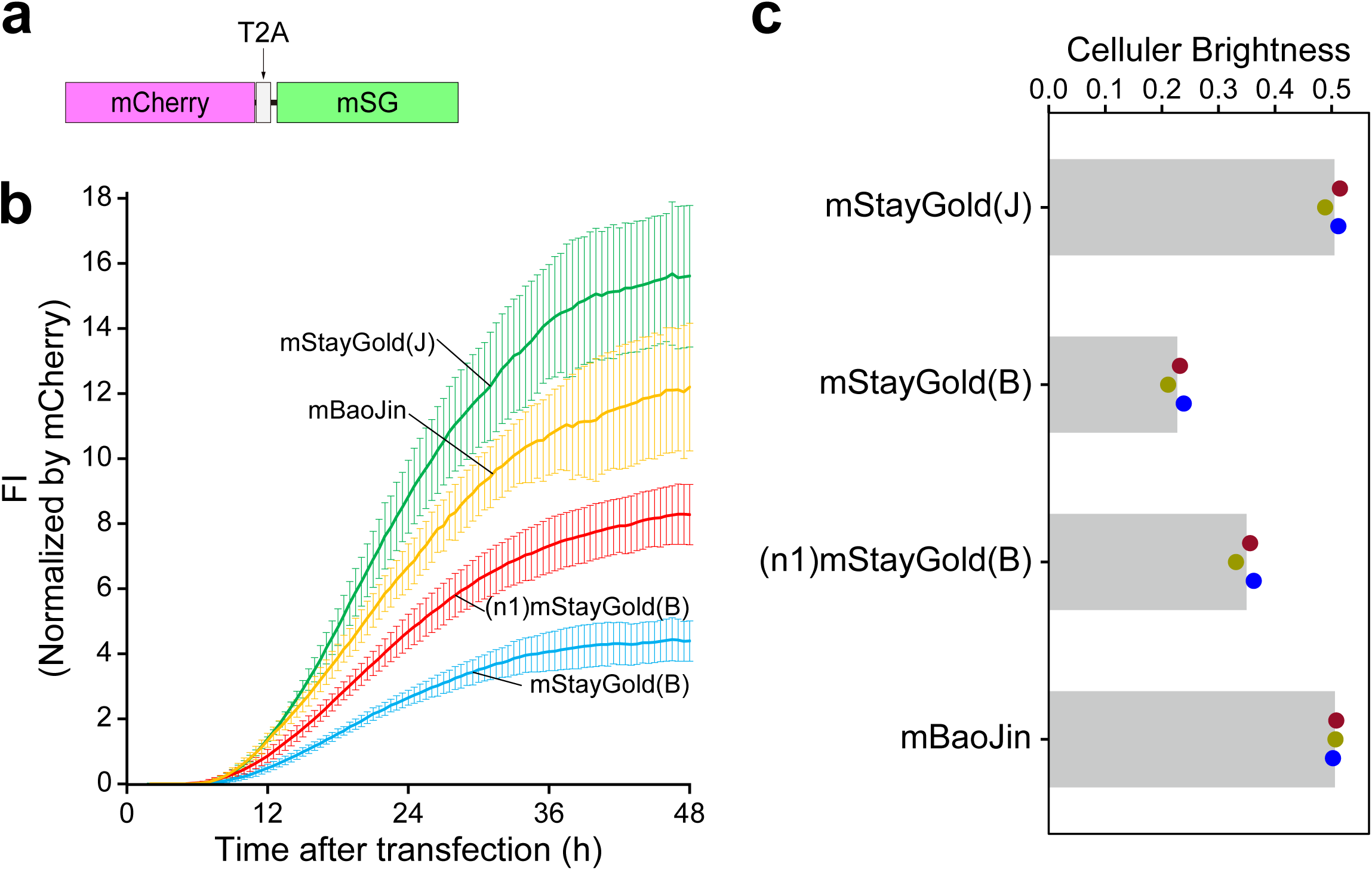
| Brightness of StayGold monomers in live HeLa cells. **a**, Cotranslation of mSG with mCherry using the bicistronic coexpression system. Transfection was performed with pCSII-EF/mCherry-T2A-mSG. mSG: StayGold monomer. **b**, Fluorescence development after transfection. FI: fluorescence intensity. Side-by-side comparison of four variants was made for their chromophore maturation using an automated time-lapse imaging system that accommodates a twelve-well plate. The green fluorescence was corrected for the mCherry fluorescence (*t* = 48 h). Data points are shown as means ± s.d. (*n* = 3 independent experiments). Traces are indicated in green for mStayGold(J), aqua for mStayGold(B), red for (n1)mStayGold, and orange for mBaoJin. **c**, Cellular brightness 48 h after transfection. Images were acquired after 48 h incubation. The green fluorescence was corrected for the mCherry fluorescence. Transfection was repeated three times; plotted dots of individual experiments are colored crimson, olive, and blue; mean values are shown by gray bars. **b, c**, The green fluorescence was first corrected for the respective spectral throughputs (Supplementary Fig. 3).

pCSII-EF/mCherry-T2A-mSG was transfected into HeLa cells. We first performed long-term, time-lapse imaging experiments by using a fully automated imaging system (SARTORIUS, Incucyte SX5) to monitor the fluorescence development after transfection (Fig. 3b). We confirmed that the fluorescence intensities of all four mSGs were gradually increasing at 48 h toward the plateau levels (Supplementary Note 2). We next determined the cellular brightness that reflected the chromophore maturation yield inside the cell at a specific time point (48 h after transfection) (Fig. 3c). mStayGold(J) and mBaoJin showed the same cellular brightness; (n1)mStayGold(B) and mStayGold(B) showed 45% and 69% brightness, respectively, compared with mStayGold(J) or mBaoJin.

### Photostability

We expressed the four mSGs in HeLa cells as fusions to histone 2B (H2B) to be immobilized on chromatin structures inside the nucleus (Fig. 4a). Twenty-four hours after transfection, we comprehensively examined all culture dishes and found that the relative brightness of the four constructs inside the nucleus was the same as observed in Figure 3 (Supplementary Fig. 4). Live cell samples were exposed to continuous illumination under an unattenuated light-emitting diode (LED) lamp (Fig. 4b). The photobleaching curves were normalized using the standard method^7^ that considered the molecular brightness of each variant at 483 nm (Supplementary Note 3) and the irradiance (8.66 W cm^−2^) (Fig. 4c). The time required for photobleaching from an initial emission rate of 1,000 to 500 photons s^−1^ molecule^−1^ (*t*_1/2_) of each variant is presented in Table 1. mStayGold(J), mStayGold(B), and (n1)mStayGold(B) showed approximately the same photostability. Compared with these three variants, mBaoJin was approximately five times less photostable.

**Figure 4.**
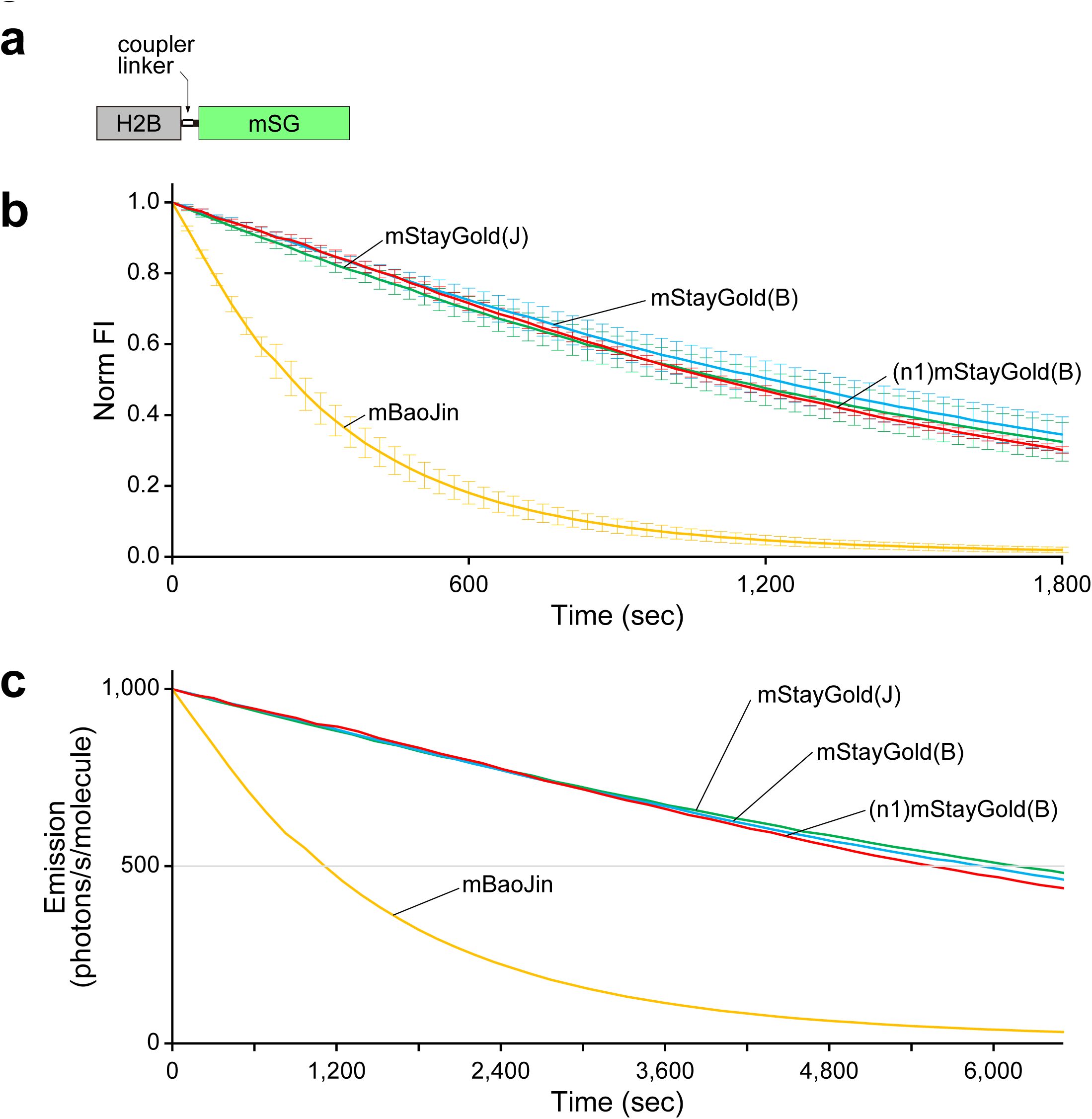
| Photostability of StayGold monomers in live HeLa cells. a, Domain structure of H2B-FP. H2B: histone 2B. mSG: StayGold monomer. b, Intensity-normalized curves. FI: fluorescence intensity. c, Plot of intensity vs. normalized total exposure time, with an initial emission rate of 1,000 photons s^−1^ molecule^−1^. Illumination intensity, 8.66 W cm^−2^. b, c, The curves are representative of three repetitions (*n* = 3 independent experiments). Traces are indicated in green for mStayGold(J), aqua for mStayGold(B), red for (n1)mStayGold, and orange for mBaoJin. Error bars indicate s.d. (b). The statistical values of *t*_1/2_ (time for photobleaching from an initial emission rate of 1,000 photons s^−1^ molecule^−1^ down to 500 photons s^−1^ molecule^−1^) are shown in Table 1.

### Multifaceted analysis using OSER cell samples

We focused on the organized smooth ER (OSER) assay^8^ as a typical example of molecular fusion and membrane targeting applications. In this assay, an FP is fused to the C-terminus of CytERM for ER membrane targeting (Fig. 5). We designed a multifaceted approach based on the OSER assay system to evaluate the practical brightness, photostability, and dispersibility of an FP. First, to quantify the expression level of each mSG, we inserted the most established tag (myc) between CytERM and mSG (Fig. 5a). After fixation with 4% paraformaldehyde (PFA) for 10 min, immunostaining using an anti-myc antibody conjugated with a red fluorophore enabled precise fusion protein quantification. This approach, although laborious, should provide more reliable information on the expression level than simple tagging with a reference red-emitting FP, the maturation of which could vary depending on the situation. Second, this experimental design enabled us to evaluate the sensitivity of each mSG to chemical fixation. Third, we used live cell images to measure OSER scores. Finally, we performed photobleaching experiments by using OSER cell samples.

**Figure 5.**
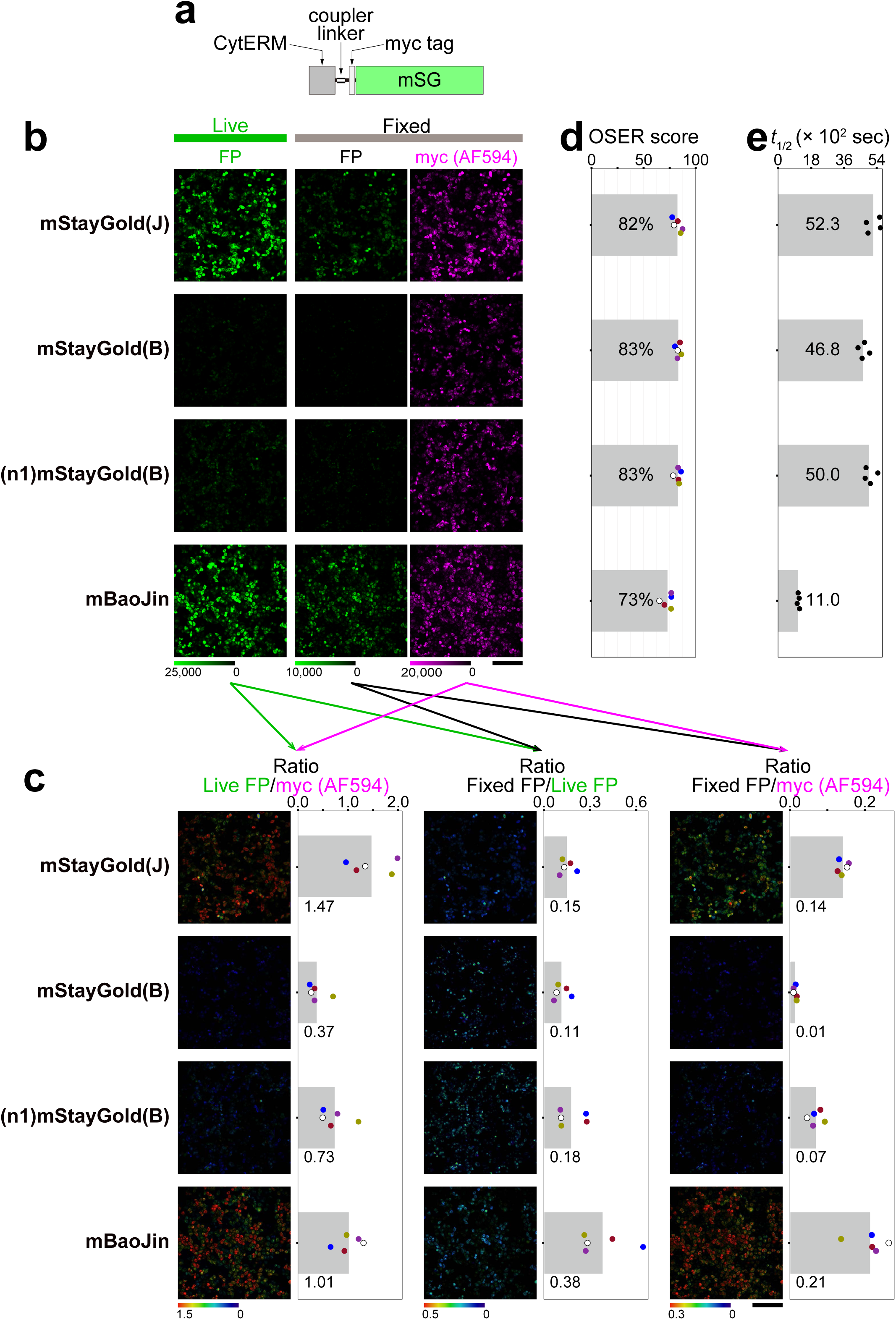
| Multifaceted analysis of four StayGold monomers fused to CytERM and targeted to the ER membrane. **a**, Domain structure of CytERM=myc-mSG. “=” denotes “Coupler linker,” a triple repeat of the amino acid linker: Gly-Gly-Gly-Gly-Ser. mSG: StayGold monomer. **b**, Green fluorescence images before (Live FP) and after (Fixed FP) fixation. Immunofluorescence (Alexa Fluor 594) images against the myc tag (Fixed myc (AF594)) (Supplementary Note 7). Color scales indicate the lowest and highest intensities of the fluorescence images. **c**, Ratio images and statistical values of Live FP/myc (AF594) (left), Fixed FP/Live FP (middle), and Fixed FP/myc (AF594) (right). Color bars indicate the lowest and highest ratio values of the images. The green fluorescence was first corrected for the respective spectral throughputs (Supplementary Fig. 3). For each variant, the ratio value is dot plotted, and the means are shown by gray bars with numerical values. **b, c**, Scale bars, 200 μm. **d**, OSER assay for dispersibility of variants. For each variant, the percentage of cells scored without visible whorl structures is dot plotted, and the mean scores are shown by gray bars with numerical values. **b–d**, Transfection was repeated five times. Plotted dots of individual experiments are colored blue, crimson, olive, violet, and white in **c** and **d**. Images of one representative experiment (white) are shown in **b** and **c**; this experiment gave the median values of the Live FP/myc (AF594) statistics. **e**, The statistical values of *t*_1/2_ (time for photobleaching from an initial emission rate of 1,000 photons s^−1^ molecule^−1^ down to 500 photons s^−1^ molecule^−1^) are plotted (*n* = 4 independent experiments). See Supplementary Fig. 5. **b–e**, from top to bottom: mStayGold(J), mStayGold(B), (n1)mStayGold(B), and mBaoJin.

Figures 5b–d show an assembly of data from five independent experiments that directly compared CytERM-mStayGold(J), CytERM-mStayGold(B), CytERM-(n1)mStayGold(B), and CytERM-mBaoJin. As revealed by immunostaining (Fig. 5b, Fixed, myc (AF594)), in one experiment, similar expression levels of the four constructs were obtained. However, bright fluorescent signals were observed live from mStayGold(J) and mBaoJin but not from mStayGold(B) or (n1)mStayGold(B) (Fig. 5b, Live, FP). The ratio (Live FP/myc (AF594)) images and the statistical values (Fig. 5c, left) indicate that the brightest per host protein (CytERM) was mStayGold(J), followed by mBaoJin, (n1)mStayGold(B), and mStayGold(B). The cellular brightness measured on the ER membrane was lowered to 25%, 50%, or 69% with mStayGold(B), (n1)mStayGold(B), or mBaoJin, respectively, compared with that of mStayGold(J) (Table 1). We again note that the addition of n1 improved the maturation of mStayGold(B) in the CytERM fusion.

Statistical ratioing with Fixed FP signals as numerators revealed that mBaoJin was the most resistant to fixation among the four mSGs (Fixed FP/Live FP and Fixed FP/myc (AF594)) (Fig. 5c, middle and right, respectively). In addition, live images of all five experiments were used to perform the OSER assay; CytERM-mStayGold(J), CytERM-mStayGold(B), CytERM-(n1)mStayGold(B), and CytERM-mBaoJin exhibited reasonably high dispersibility, with OSER scores of 82%, 83%, 83%, and 73%, respectively (Fig. 5d). Apart from the above, three independent transfections were performed using the four CytERM-mSG cDNAs to study their photostability. Photobleaching curves (Supplementary Fig. 5) very similar to those shown in Figure 4 were obtained; the *t*_1/2_ values are presented in Figure 5e.

### Fluoppi for analysis of brightness as well as dispersibility

To evaluate the dispersibility of StayGold variants, we used a Fluoppi-based assay^2,9^ in addition to the OSER assay. We expressed the four mSGs in HeLa cells as fusions to the Phox and Bem1p (PB1) domain of p62/SQSTM1 (Supplementary Fig. 6a). We obtained high Fluoppi scores with all four of the mSG constructs (Supplementary Fig. 6b). After side-by-side transfections, however, we observed a considerable difference in brightness between two groups: mStayGold(J) or mBaoJin produced fluorescence approximately one order of magnitude brighter than that of mStayGold(B) or (n1)mStayGold(B) (Supplementary Fig. 6c). Although this comparison was not as quantitative as the ratiometric approaches that employed internal references such as the mCherry (Fig. 3) and myc tag (Fig. 5), we reproducibly observed such a difference in brightness, which was statistically analyzed (Supplementary Fig. 6d).

## Discussion

In the early phase of the directed evolution toward StayGold monomers, we were also interested in the glutamate residue at position 138 (Glu^138^). We mutated this amino acid to all the others in His6-(n1)oxStayGold (Supplementary Note 4) and found that most mutations, including E138D, rendered the bacterial colonies very dim. Because we considered brightness in bacteria to be a requisite, we did not characterize these mutants further. As we suspected that Glu^138^ might be critical for the photostability of this FP, we were very impressed by how the E138D mutation was able to promote the monomerization of StayGold without affecting its photostability^4^. Using mStayGold(B), we verified that this StayGold monomer showed high photostability (Fig. 4) and molecular brightness (Table 1). We expect that mStayGold(B) will contribute to single-molecule measurements that require a low density of fluorescently labeled proteins. However, we argue that mStayGold(B) is practically dim on the following grounds. According to our quantitative measurement (cotranslation with mCherry) of cellular brightness, which is usually measured in the cytosolic and nuclear compartments of cultured mammalian cells, mStayGold(B) showed approximately 45% brightness compared with mStayGold(J) (Fig. 3). Owing to the poor maturation probably when targeted to the ER membrane, the brightness per CytERM fusion of mStayGold(B) was further lowered to approximately 25% relative to that of mStayGold(J) (Fig. 5). Because nonfluorescent fusion proteins occupy designated spaces inside the cell without being detected, every effort should be made to minimize their fraction. Relative darkness of mStayGold(B) was also noticed when it was expressed as a fusion to His6 in bacterial cells (Fig. 2) and as a fusion to PB1 in HeLa cells (Supplementary Fig. 6). In all of these cases (Figs. 2, 3, 5 and Supplementary Figs. 4, 6), the addition of the n1 adaptor at the N-terminus of mStayGold(B) was effective for brightness recovery, albeit partially given the brightness of mStayGold(J). In contrast to C-terminal tagging of POIs, which was featured in the present study, the insertion of an appropriate adaptor, such as c4 and PT, at the C-terminus of these StayGold monomers is recommended in reverse fusions^2^ (Supplementary Fig. 7).

Disrupting the dimerization of StayGold would likely result in the loss of either brightness or photostability. High-throughput screening is easy to perform for bright FPs but not for photostable FPs. By adopting a low-throughput approach to select photostable FPs during the directed evolution, however, we were able to obtain mStayGold(J), which is both photostable and bright. By contrast, mBaoJin appears to be a product of numerous rounds of directed evolution that prioritized only brightness^5^. It is thus reasonable that mBaoJin is less photostable than the original StayGold. Our data indicate that mBaoJin exhibits approximately 20% photostability in living cells compared with mStayGold(J) or mStayGold(B) (Figs. 4 and 5e). We measured FP photobleaching rates in living cells according to Dr. Shaner’s opinion^10^ and obtained similar results from cells where mSGs were localized to the nucleus (Fig. 4) and ER membrane (Fig. 5). Despite the reduced photostability, however, mBaoJin is still more photostable than common bright green-emitting FPs, such as mNeonGreen (Supplementary Note 5), and should contribute to improving super-resolution long-term live cell imaging^5^. With extensions at both the N– and C-termini, furthermore, mBaoJin appears to be an all-purpose tag (Supplementary Note 6). We are instead strongly interested in mBaoJin’s extreme resistance to chemicals such as fixatives and guanidium hydrochloride^5^. This discovery is attributable to the versatile approaches taken by the research group that developed mBaoJin. mBaoJin is expected to empower the technologies of tissue clearing/expansion^11^ and correlative light and electron microscopy^12^.

In summary, we have a high opinion of both the strategies of the two research groups that rival ours and the results derived from them. However, we think that both mStayGold(B) and mBaoJin are partially developed products in the proper context of StayGold monomerization. They remind us of the critical tradeoff between the photostability and brightness of FPs that we discussed in depth in our original paper on StayGold^1^. Lastly, because we do not yet understand the structural basis of the high photostability of StayGold, room still exists for further improvements in the three basic properties—photostability, brightness, and dispersibility—of the monomers, including mStayGold(J).

*Note added in proof* (February 27): During the submission of this preprint manuscript, we came across the excellent work of Drs. Piatkevich and Subach and their colleagues on mBaoJin in Nature Methods (https://doi.org/10.1038/s41592-024-02203-y). It appears that mBaoJin is a versatile FP; it is less photostable but more stable against many environmental factors than mStayGold(J) and mStayGold(B). We are aware of the many issues surrounding the variation in physicochemical properties, especially photostability, of StayGold derivatives in solution vs. living cells, before vs. after chemical fixation, and in the presence vs. the absence of N– or C-terminal extensions. These issues need further clarification, and we would like to reserve related discussions for our forthcoming publications. In the present study, we have made a simple but reasonable effort to address the issue of how to expand a triangle that has the photostability, brightness, and dispersibility of StayGold monomers at its three vertices. We hope that StayGold technology will be further refined and diversified for many biological applications.

## Acknowledgments

The authors thank K. Higuchi, Y. Ue, and Y. Chiba at the RIKEN CBS-Evident Collaboration Center for technical assistance with LSCM; Common Use Equipment in the Support Unit for Bio-Material Analysis, Research Resource Division, RIKEN CBS for technical assistance with Incucyte SX5; Dr. I. Imayoshi at Kyoto University and Dr. R. Kageyama at RIKEN CBS for continuous support; Dr. T. Fujiwara at Kyoto University, Dr. Y. Okada at the University of Tokyo and Dr. H. Kurokawa at RIKEN CBS for valuable advice. This work was supported in part by Grant-in-Aid for Scientific Research (S) (21H05041), Grant-in-Aid for Innovative Areas: Resonance Bio (15H05948), Marine Biomass Innovation Project (NFRFT-2020-00452), the Brain Mapping by Integrated Neurotechnologies for Disease Studies from AMED (Brain/MINDS, JP15dm0207001).

## Author Contributions

S.S. and R.A. analyzed dispersibility and cellular brightness of mSGs. R.A. analyzed photobleaching of mSGs. M.H. analyzed spectroscopic properties of mSGs. Y.N. analyzed brightness of mSGs in bacteria. S.S., R.A., M.H. and Y.N. carried out gene construction. A.M. and M.S. prepared figures. A.M. designed the study and wrote the manuscript.

## Methods

### Protein purification

Recombinant proteins with a polyhistidine tag at the N-terminus were expressed in *Escherichia coli* [JM109 (DE3)]. Transformed *E. coli* were incubated in a Luria–Bertani (LB) medium containing 0.1 mg mL^−1^ ampicillin at room temperature (RT) with gentle shaking for several days. Protein purification by Ni^2+^ affinity chromatography was performed as described previously^13^.

### In vitro spectroscopy

Absorption spectra were acquired using a spectrophotometer (U-2910, Hitachi). Fluorescence excitation and emission spectra were acquired using a fluorescence spectrophotometer (F-2500, Hitachi). Absolute fluorescence quantum yields were measured using an absolute photoluminescence quantum yield spectrometer (C9920-02, Hamamatsu Photonics). The solution for spectroscopy contained 50 mM HEPES (KOH) pH 7.4 and 150 mM KCl. Protein concentrations were measured using a Protein Assay Dye Reagent Concentrate kit (#5000006, Bio-Rad) with bovine serum albumin (BSA) as the standard. See Figs. 2b–d.

### FP maturation in bacterial cells

A homemade fluorescence analyzing system consisting of a Xenon light source MAX-302 (Asahi Spectra), an excitation filter (465–495 nm) (480AF30, Omega Optical), an emission filter (530–550 nm) (PB0540/020, Asahi Spectra), and a sCMOS camera ZYLA-5.5-USB3 (Andor) was used for time-lapse imaging of transformed *E. coli* colonies that expressed mStayGold(J), mStayGold(B), (n1)mStayGold(B) or mBaoJin. The whole system was controlled by MetaMorph software (Molecular Devices). Multiple colonies were made for each FP by spotting 1.5-μL drops of transformed competent JM109(DE3) cell suspension on an LB agar plate with 100 µg mL^−1^ ampicillin. After 2 h incubation at 37 °C, the plate was placed in a stage-top incubation chamber (IBC, Tokai Hit) kept at 37 °C and time-lapse imaging was immediately started. Images were analyzed using ImageJ (National Institutes of Health). See Fig. 2e and Supplementary Fig. 3.

### Gene construction for bicistronic expression in mammalian cells

The StayGold monomer (mSG) gene was amplified using primers containing 5’-*Bam*HI and 3’-*Xba*I sites, and the restricted product was cloned in frame into the *Bam*HI/*Xba*I sites of pBS/mCherry-T2A (ref. 2) to generate pBS/mCherry-T2A-mSG. Lastly, *Xho*I/*Xba*I fragments encoding mCherry-T2A-mSG were subcloned into pCSII-EF to generate pCSII-EF/mCherry-T2A-mSG plasmids. See Fig. 3a.

### FP maturation in mammalian cells

HeLa cells seeded into 12-well plates (353043, CORNING) and maintained in growth medium (Dulbecco’s Modified Eagle Medium (DMEM) low glucose, supplemented with 10% fetal bovine serum). On the following day, cells were transfected with 0.5 µg of pCSII-EF/mCherry-T2A-mSG per well using 1 µL of Lipofectamine 2000 (52887, Thermo Fisher). After one-h incubation with the transfection complexes, the medium was replaced with fresh phenol-red free DMEM (044-33555, Fuji Film) supplemented with 10% FBS and GlutaMax (35050061, Thermo Fisher). One hour after the removal of the transfection complexes, cells were subjected to long-term, time-lapse imaging using a fully automated imaging system (SARTORIUS, Incucyte SX5) that was maintained at 37 °C in a 5% CO_2_ environment in an incubator (Thermo Fisher, Forma Steri-Cycle i250). Fluorescence and phase contrast images (4 images per well per channel) were acquired every 30 min using a 10× objective lens and the G/O/NIR Filter Set. StayGold monomers were observed using the G channel (excitation: 453–485 nm, emission: 494–533 nm) (Supplementary Fig. 3). mCherry was observed using the O channel (excitation: 546–568 nm, emission: 576–639 nm). FP signals were defined as pixels having signal values exceeding 5 times the standard deviation above the mean fluorescence intensity of the first images.

Because the fluorescence development of StayGold monomers preceded that of mCherry, the green/red ratios increased abruptly in the early phase. Therefore, each signal of a StayGold monomer was divided by the respective mCherry signal at 48 h. See Fig. 3b.

### Cellular brightness (normalized against mCherry)

HeLa cells were seeded into 24-well glass-bottom plates (5826-024, IWAKI) and maintained in growth medium (Dulbecco’s Modified Eagle Medium low glucose, supplemented with 10% fetal bovine serum). On the following day, cells were transfected with 0.5 μg of pCSII-EF/mCherry-T2A-mSG per well using 1 µL of Lipofectamine 2000 (52887, Thermo Fisher). The fluorescence of both mSG and mCherry was distributed evenly throughout the cell. Forty-eight hours after transfection, cells were incubated in Hanks’ Balanced Salt Solution (HBSS, 14025, Thermo Fischer Scientific) containing 15 mM HEPES-NaOH (pH 7.4) and imaged on an inverted microscope (IX-83, Evident) equipped with an LED light bulb (X-Cite XYLIS, Excelitas Technologies), an objective lens (UPlanXApo 4×/0.16 NA, Evident), and a scientific CMOS camera (ORCA-Fusion, Hamamatsu Photonics). StayGold monomers were observed using a filter cube (U-FBNA, Evident), which is composed of an excitation filter (470–495 nm), a dichroic mirror (505LP), and an emission filter (510–550 nm) (Supplementary Fig. 3). mCherry was observed using a filter cube (U-FMCHE, Evident), which is composed of an excitation filter (565–585 nm), a dichroic mirror (595LP), and an emission filter (600–690 nm). The StayGold monomer fluorescence was corrected for the mCherry fluorescence. To correct the StayGold monomer fluorescence by the mCherry fluorescence, the StayGold monomer image was divided by the mCherry image. A mask was created by applying the ‘MinError’ algorithm to the mCherry image using Fiji (version 2.14.0/1.54f). Saturated pixels in either the StayGold monomer or the mCherry channel were subsequently removed from the mask. This mask was then applied to the ratio image to calculate the mean of the corrected brightness of the StayGold monomer. See Fig. 3c.

### Gene construction (nuclear targeting)

The mouse histone 2B (H2B) gene (Fantom3) was amplified using primers containing 5’-*Xho*I and 3’-*Hin*dIII sites, and the restricted product was cloned into the *Xho*I/*Hin*dIII sites of pBS Coupler 1 (ref. 14) to generate pBS Coupler 1/H2B. In addition, the StayGold monomer (mSG) gene was amplified using primers containing 5’-*Bam*HI and 3’-*Xba*I sites, and the restricted product was cloned in frame into the *Bam*HI/*Xba*I sites of pBS Coupler 1/H2B to generate pBS Coupler 1/H2B=mSG. Lastly, *Xho*I/*Xba*I fragments encoding H2B=mSGs were subcloned into pCSII-EF for transfection.

### Gene construction (ER membrane targeting)

A *Kpn*I/*Bam*HI gene fragment encoding a Coupler linker and a myc tag (EQKLISEEDL) was created by PCR. The gene fragment was cloned into the *Kpn*I/*Bam*HI sites of pcDNA3/F-tractin=mStayGold(J) (ref. 2) to generate pcDNA3/F-tractin=myc-mStayGold(J). Then, the CytERM gene derived from pcDNA3/CytERM-FP (ref. 2) was substituted for the F-tractin gene at the *Hin*dIII/*Kpn*I sites to generate pcDNA3/CytERM=myc-mStayGold(J). Lastly, the mStayGold(B), (n1)mStayGold(B), or mBaoJin gene was substituted for the mStayGold(J) gene at the *Bam*HI/*Xho*I sites to generate pcDNA3/CytERM=myc-mStayGold(B), pcDNA3/CytERM=myc-(n1)mStayGold(B), or pcDNA3/CytERM=myc-mBaoJin, respectively.

### WF photobleaching

HeLa cells were seeded on 35-mm glass-bottom dishes and maintained in growth medium (Dulbecco’s Modified Eagle Medium low glucose, supplemented with 10% fetal bovine serum). On the following day, cells were transfected with 0.5 μg of plasmid DNA per well using 2.5 µL of 1 mg mL^−1^ polyethylenimine (PEI) (Linear, MW 25,000, Polysciences Inc.). Twenty-four hours after transfection, living cells expressing H2B=mSG or CytERM=myc-mSG were incubated in HBSS containing 15 mM HEPES-NaOH (pH 7.4) and imaged on an inverted microscope (IX-83, Evident) equipped with an LED light bulb (X-Cite XYLIS, Excelitas Technologies), a 60× objective lens (UPlanSApo 60×/1.35 NA), a scientific CMOS camera (ORCA-Fusion, Hamamatsu Photonics), and a filter cube (U-FBNA, Evident), which is composed of an excitation filter (470–495 nm), a dichroic mirror (505LP), and an emission filter (510–550 nm) (Supplementary Fig. 3). Cells were excited continuously; the central wavelength of the excitation passband was approximately 483 nm. Image acquisition was performed every 30 sec with a short exposure time (20 ms). The data were analyzed using Excel (2019). The fluorescence intensity at t = 0 was normalized to 1,000 photons s^−1^ molecule^−1^, and the time axis was adjusted according to the standard method^7^. The power of excitation light (W) above the objective at the focal plane was measured using a microscope slide power meter sensor (S170C; Thorlabs, Newton, NJ) and an optical power and energy meter (PM100D; Thorlabs, Newton, NJ). The power was divided by the area of the illumination field (cm^2^) to obtain the irradiance. In all cases, the illuminator (collimator lens) was adjusted to achieve Köhler illumination. A color acrylic plate (Tokyu Hands, Japan) was placed at the focal plane to evaluate illumination uniformity on a sCMOS image. See Figs. 4 and 5e.

### OSER assay

HeLa cells were seeded into 24-well glass-bottom plates (5826-024, IWAKI) and maintained in growth medium (Dulbecco’s Modified Eagle Medium low glucose, supplemented with 10% fetal bovine serum). On the following day, cells were transfected with 0.5 μg of pcDNA3/CytERM=myc-mSG per well using 1 µL of Lipofectamine 2000 (52887, Thermo Fisher). Twenty-four hours after transfection, cells were incubated in HBSS containing 15 mM HEPES-NaOH (pH 7.4) and imaged on an inverted microscope (IX-83, Evident) equipped with a 20× objective lens (UPlanXApo 20×/0.8 NA, Evident), a camera (ORCA-Fusion, Hamamatsu Photonics), and a filter cube (U-FBNA, Evident). At an *xy* position, nine images were serially acquired with a *z*-step size of 0.59 µm, from which an in-focus image was mathematically generated by the extended focus imaging (EFI) function of the cellSens Dimension (Evident) software (Version 3.2). The number of transfected cells showing whorl structures was counted. In addition, the number of transfected cells avoiding whorl formation was counted. See Fig. 5d. The images were also treated as ‘Live FP’ (Figs. 5b, c).

### Cellular brightness (normalized against myc)

After the OSER assay, cells were fixed with 4% paraformaldehyde (PFA)/phosphate-buffered saline (PBS) at RT for 10 min. After being washed in PBS three times, cells were incubated in blocking solution (PBS containing 3% BSA and 0.1% Triton X-100) at RT for 60 min. The cells were reacted with Alexa Fluor 594-conjugated primary antibody (anti-myc-tag mAb, Medical Biological Laboratory, #M047-A59, clone: PL14) in blocking solution at RT for 60 min. After being washed in PBS containing 0.1% Triton X-100 three times, cells were incubated in HBSS containing 15 mM HEPES-NaOH (pH 7.4) and imaged on an inverted microscope (IX-83, Evident) equipped with a 20× objective lens (UPlanXApo 20×/0.8 NA, Evident) and a camera (ORCA-Fusion, Hamamatsu Photonics). StayGold monomers were observed using a filter cube (U-FBNA, Evident), which is composed of an excitation filter (470–495 nm), a dichroic mirror (505LP), and an emission filter (510–550 nm) (Supplementary Fig. 3). Alexa Fluor 594 was observed using a filter cube (U-FMCHE, Evident), which is composed of an excitation filter (565–585 nm), a dichroic mirror (595LP), and an emission filter (600–690 nm). These green and red fluorescence images were treated as ‘Fixed FP’ and ‘myc (AF594)’, respectively (Figs. 5b, c). For correction of fixation-induced change in morphology, alignment was performed between ‘Live’ and ‘Fixed’ images using the Fiji MultiStackReg plug-in with the Rigid Body algorithm (Fiji version: 2.14.0/1.54f).

### Fluoppi assay

The StayGold monomer gene was amplified using primers containing 5’-*Bam*HI and 3’-*Eco*RI sites, and the restricted product was cloned into the *Bam*HI/*Eco*RI sites of pAsh-MCL (Medical Biological Laboratory, Japan) to generate a plasmid DNA for expression of PB1-mSG. HeLa cells were seeded on a 35-mm glass-bottom dish and maintained in growth medium (Dulbecco’s Modified Eagle Medium low glucose, supplemented with 10% fetal bovine serum). Twenty-four hours after transfection with 0.5 μg of plasmid DNA per well using 2.5 µL of 1 mg mL^−1^ polyethylenimine (PEI) (Linear, MW 25,000, Polysciences Inc.), cells were incubated in HBSS containing 10 mM HEPES-NaOH (pH 7.4) and imaged on an inverted microscope (IX-83, Evident) equipped with a 20× objective lens (UPlanXApo 20×/0.8 NA, Evident) and a camera (ORCA-Fusion, Hamamatsu Photonics). The filter cube used was U-FBNA (Evident). The numbers of transfected cells showing and avoiding puncta were counted individually. See Supplementary Fig. 5b.

### Cellular brightness (without normalization)

HeLa cells were seeded on 35-mm glass-bottom dishes and maintained in growth medium (Dulbecco’s Modified Eagle Medium low glucose, supplemented with 10% fetal bovine serum). On the following day, transfection was carried out using 0.5 μg of plasmid DNA and 2.5 µL of 1 mg mL^−1^ polyethylenimine (PEI) (Linear, MW 25,000, Polysciences Inc.). Twenty-four hours after transfection, cells were incubated in HBSS containing 15 mM HEPES-NaOH (pH 7.4) and imaged on an inverted microscope (IX-83, Evident) equipped with an LED light bulb (X-Cite XYLIS, Excelitas Technologies), a 20× objective lens (UPlanXApo 20×/0.8 NA, Evident), a scientific CMOS camera (ORCA-Fusion, Hamamatsu Photonics), and a filter cube (U-FBNA, Evident), which is composed of an excitation filter (470–495 nm), a dichroic mirror (505LP), and an emission filter (510–550 nm) (Supplementary Fig. 3). See Supplementary Figs. 4b, 6d.

## Data availability

The accession numbers in the DDBJ/GenBank databases are LC797985 for mStayGold(B), LC797986 for (n1)mStayGold(B), and LC797987 for mBaoJin that has a gene with mammalian-preferred codons. All data generated in this study are available through the RIKEN Research Data & copyrighted-work Management System (https://dmsgrdm.riken.jp/shwmu/). Plasmid DNAs containing StayGold monomers are available from the RIKEN Bio-Resource Center (BRC) (http://en.brc.riken.jp) under a material transfer agreement with RIKEN.

