## Supplementary Information for "Comparison of monomeric variants of StayGold"

### Table of Contents

**Supplementary Notes 1 - 7**

**Supplementary Figures 1 - 7**

**Supplementary References**

### Supplementary Notes

#### Supplementary Note 1
This monomeric variant of StayGold appears by the name of StayGold-E138D in the FPbase database and mStayGold (E138D) at addgene.

#### Supplementary Note 2
Here we wanted to confirm similar temporal profiles of the normalized fluorescence intensities among the four StayGold monomers. For example, an exception is seen with mNeonGreen. The normalized fluorescence of mNeonGreen peaked at approximately 30 h and gradually decreased thereafter, indicating degradation of the β-barrel of this FP inside the cell[1].

#### Supplementary Note 3
An excitation bandpass filter (470–495 nm) was used for photobleaching StayGold monomers. The central wavelength of the excitation passband was approximately 483 nm.

#### Supplementary Note 4
oxStayGold contains three mutations (H169Y, C174I, and C208I, see Figure 1) relative to StayGold and remains a dimer[1,2].

#### Supplementary Note 5
Most bright green-emitting FPs, such as mNeonGreen, show only a few percent photostability compared with mStayGold(J) according to our previous studies[1,2].

#### Supplementary Note 6
In our tool palette, mStayGold2, which has n1 and PT adaptors at the N- and C-termini, respectively, is a similar all-purpose tag[1]

#### Supplementary Note 7
We confirmed that no signal was observed for myc in non-transfected cells.

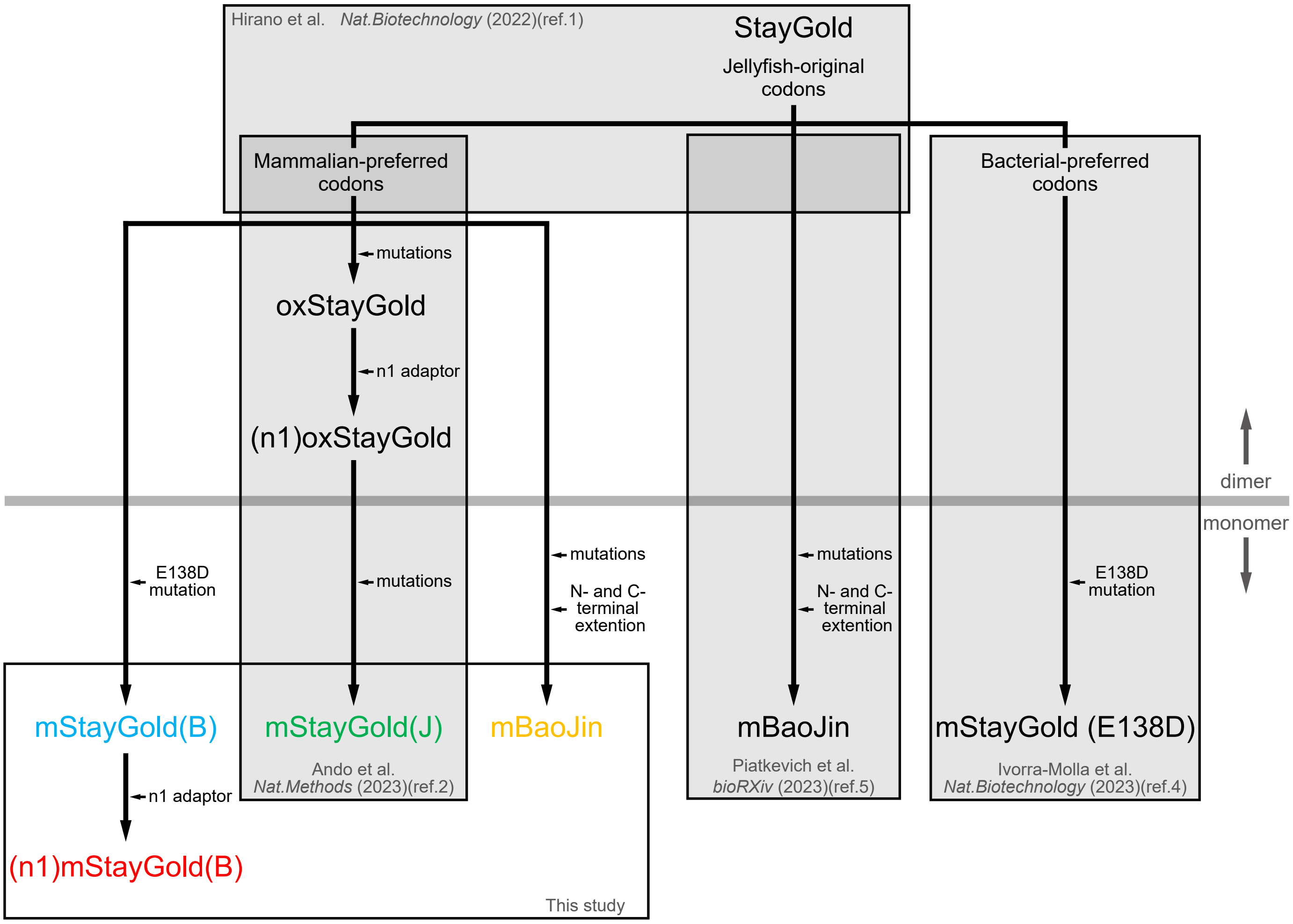


**Supplementary Fig. 1**
Schematic illustrating how mStayGold(J), mStayGold(B), (n1)mStayGold(B), and mBaoJin were prepared from StayGold. Variants featured in each publication are boxed.

### mStayGold (E138D) (bacterial-preferred codons) vs. mStayGold(B) (mammalian-preferred codons)

```
mStayGold(B)     M   A   S   T   P   F   K   F   Q   L       K   G   T   I   N   G   K   S   F   T       V   E   G   E   G   E   G   N   S   H
  (bacterial)   ATG GCC AGC ACG CCG TTT AAG TTC CAG TTA   AAG GGG ACA ATT AAC GGG AAA AGC TTT ACA   GTC GAG GGT GAA GGG GAA GGG AAC TCG CAT
  (mammalian)   ATG GCC AGC ACC CCC TTC AAG TTC CAG CTG   AAG GGC ACC ATC AAC GGC AAG AGC TTC ACC   GTG GAA GGC GAG GGC GAG GGC AAT AGC CAC
                *** *** *** **  **  **  *** *** ***  *    *** **  **  **  *** **  **  *** **  **    **  **  **  **  **  **  **  **      **

mStayGold(B)     E   G   S   H   K   G   K   Y   V   C       T   S   G   K   L   P   M   S   W   A       A   L   G   T   S   F   G   Y   G   M
  (bacterial)   GAA GGT AGT CAT AAA GGT AAG TAT GTG TGT   ACG AGC GGA AAG TTG CCG ATG TCT TGG GCG   GCT CTT GGT ACT TCG TTT GGG TAC GGA ATG
  (mammalian)   GAG GGC AGC CAC AAA GGC AAG TAC GTG TGC   ACC AGC GGC AAA CTG CCA ATG TCT TGG GCC   GCC CTG GGA ACT AGC TTC GGC TAT GGC ATG
                **  **  **  **  *** **  *** **  *** **    **  *** **  **   ** **  *** *** *** **    **  **  **  ***     **  **  **  **  ***

mStayGold(B)     K   Y   Y   T   K   Y   P   S   G   L       K   N   W   F   H   E   V   M   P   E       G   F   T   Y   D   R   H   I   Q   Y
  (bacterial)   AAA TAC TAT ACA AAA TAC CCG TCG GGC CTG   AAA AAT TGG TTC CAT GAG GTG ATG CCC GAG   GGG TTC ACG TAT GAT CGC CAC ATC CAG TAC
  (mammalian)   AAG TAC TAC ACC AAG TAC CCC AGC GGC CTG   AAG AAC TGG TTC CAC GAG GTG ATG CCC GAG   GGC TTC ACC TAC GAC AGA CAC ATC CAG TAC
                **  *** **  **  **  *** **      *** ***   **  **  *** *** **  *** *** *** *** ***   **  *** **  **  **      *** *** *** ***

mStayGold(B)     K   G   D   G   S   I   H   A   K   H       Q   H   F   M   K   N   G   T   Y   H       N   I   V   E   F   T   G   Q   D   F
  (bacterial)   AAA GGT GAT GGC TCA ATC CAC GCA AAG CAC   CAA CAC TTC ATG AAG AAT GGG ACT TAT CAT   AAC ATT GTC GAG TTT ACT GGC CAG GAC TTC
  (mammalian)   AAG GGC GAC GGC AGC ATC CAC GCC AAG CAC   CAG CAC TTC ATG AAG AAC GGC ACC TAC CAC   AAC ATC GTG GAG TTC ACC GGC CAG GAC TTC
                **  **  **  ***     *** *** **  *** ***   **  *** *** *** *** **  **  **  **  **    *** **  **  *** **  **  *** *** *** ***

mStayGold(B)     K   E   N   S   P   V   L   T   G   D       M   N   V   S   L   P   N   D   V   Q       H   I   P   R   D   D   G   V   E   C
  (bacterial)   AAG GAA AAC AGC CCC GTA TTG ACA GGT GAC   ATG AAT GTG AGC TTG CCG AAC GAT GTG CAG   CAT ATT CCA CGT GAC GAC GGA GTT GAA TGT
  (mammalian)   AAG GAG AAC AGC CCC GTG CTG ACC GGC GAC   ATG AAC GTG AGC CTG CCC AAC GAC GTG CAG   CAC ATC CCC AGA GAT GAC GGC GTG GAG TGC
                *** **  *** *** *** **   ** **  **  ***   *** **  *** ***  ** **  *** **  *** ***   **  **  **   *  **  *** **  **  **  **

mStayGold(B)     P   V   T   L   L   Y   P   L   L   S       D   K   S   K   C   V   E   A   H   Q       N   T   I   C   K   P   L   H   N   Q
  (bacterial)   CCC GTC ACA TTG TTA TAC CCC TTG CTT AGC   GAT AAA TCA AAG TGT GTC GAA GCA CAT CAA   AAC ACT ATC TGC AAA CCT CTT CAT AAC CAA
  (mammalian)   CCA GTG ACC CTG CTG TAC CCT CTG CTG AGC   GAC AAG AGC AAG TGC GTG GAG GCC CAC CAG   AAC ACC ATC TGC AAG CCC CTG CAC AAT CAG
                **  **  **   **  *  *** **   ** **  ***   **  **      *** **  **  **  **  **  **    *** **  *** *** **  **  **  **  **  **

mStayGold(B)     P   A   P   D   V   P   Y   H   W   I       R   K   Q   Y   T   Q   S   K   D   D       T   E   E   R   D   H   I   C   Q   S
  (bacterial)   CCT GCG CCA GAT GTA CCG TAC CAT TGG ATT   CGC AAA CAA TAT ACA CAG TCT AAG GAT GAT   ACG GAG GAA CGC GAT CAT ATC TGT CAG TCC
  (mammalian)   CCA GCC CCC GAT GTG CCA TAC CAC TGG ATC   AGA AAG CAG TAC ACC CAG AGC AAG GAC GAC   ACC GAG GAG AGA GAC CAC ATC TGC CAG AGC
                **  **  **  *** **  **  *** **  *** **     *  **  **  **  **  ***     *** **  **    **  *** **   *  **  **  *** **  ***   *

mStayGold(B)     E   T   L   E   A   H   L   *
  (bacterial)   GAG ACG TTA GAA GCT CAT CTT TAA
  (mammalian)   GAG ACC CTG GAG GCC CAC CTG TAA
                *** **   *  **  **  **  **  ***
```

### mBaoJin (original vs. mammalian-preferred codons)

```
mBaoJin          M   V   S   K   G   E   E   E   N   M       A   S   T   P   F   K   F   Q   L   K       G   T   I   N   G   K   S   F   T   V
  (original)    ATG GTC TCA AAG GGA GAG GAG GAA AAC ATG   GCT AGT ACA CCA TTT AAA TTT CAA CTT AAA   GGA ACC ATC AAT GGC AAA TCG TTT ACC GTT
  (mammalian)   ATG GTG AGC AAG GGC GAG GAG GAA AAC ATG   GCC AGC ACC CCC TTC AAG TTC CAG CTG AAG   GGC ACC ATC AAC GGC AAG AGC TTC ACC GTG
                *** **      *** **  *** *** *** *** ***   **  **  **  **  **  **  **  **  **  **    **  *** *** **  *** **      **  *** **

mBaoJin          E   G   E   G   E   G   N   S   H   E       G   S   H   K   G   K   Y   V   C   T       S   G   K   L   P   M   S   W   A   A
  (original)    GAA GGC GAA GGT GAA GGG AAC TCA CAT GAA   GGT TCT CAT AAA GGA AAA TAT GTT TGT ACA   AGT GGA AAA CTA CCG ATG TCA TGG GCA GCA
  (mammalian)   GAA GGC GAG GGC GAG GGC AAT AGC CAC GAG   GGC AGC CAC AAA GGC AAG TAC GTG TGC ACC   AGC GGC AAA CTG CCA ATG TCT TGG GCC GCC
                *** *** **  **  **  **  **      **  **    **      **  *** **  **  **  **  **  **    **  **  *** **  **  *** **  *** **  **

mBaoJin          L   G   T   T   F   G   Y   G   M   K       Y   Y   T   K   Y   P   S   G   L   K       N   W   F   R   E   V   M   P   G   G
  (original)    CTT GGG ACA ACC TTT GGT TAT GGA ATG AAA   TAT TAT ACC AAA TAT CCT AGT GGA CTG AAG   AAC TGG TTT CGT GAA GTA ATG CCC GGA GGC
  (mammalian)   CTG GGA ACT ACC TTC GGC TAT GGC ATG AAG   TAC TAC ACC AAG TAC CCC AGC GGC CTG AAG   AAC TGG TTC CGC GAG GTG ATG CCC GGC GGC
                **  **  **  *** **  **  *** **  *** **    **  **  *** **  **  **  **  **  *** ***   *** *** **  **  **  **  *** *** **  ***

mBaoJin          F   T   Y   D   R   H   I   Q   Y   K       G   D   G   S   I   H   A   K   H   Q       H   F   M   K   N   G   T   Y   H   N
  (original)    TTT ACC TAC GAT CGT CAT ATT CAA TAT AAA   GGC GAT GGG AGT ATC CAT GCA AAA CAC CAA   CAC TTT ATG AAA AAT GGG ACT TAT CAC AAC
  (mammalian)   TTC ACC TAC GAC AGA CAC ATC CAG TAC AAG   GGC GAC GGC AGC ATC CAC GCC AAG CAC CAG   CAC TTC ATG AAG AAC GGC ACC TAC CAC AAC
                **  *** *** **   *  **  **  **  **  **    *** **  **  **  *** **  **  **  *** **    *** **  *** **  **  **  **  **  *** ***

mBaoJin          I   V   E   F   T   G   Q   D   F   K       E   N   S   P   V   L   T   G   D   M       N   V   S   L   P   N   E   V   P   Q
  (original)    ATT GTA GAA TTT ACC GGT CAG GAT TTT AAA   GAA AAT AGT CCA GTC TTA ACT GGA GAT ATG   AAT GTC TCA TTA CCG AAT GAA GTC CCA CAA
  (mammalian)   ATC GTG GAG TTC ACC GGC CAG GAC TTC AAG   GAG AAC AGC CCC GTG CTG ACC GGC GAC ATG   AAC GTG AGC CTG CCC AAC GAG GTG CCG CAG
                **  **  **  **  *** **  *** **  **  **    **  **  **  **  **   *  **  **  **  ***   **  **       *  **  **  **  **  **  **

mBaoJin          I   P   R   D   D   G   V   E   C   P       V   T   L   L   Y   P   L   L   S   D       K   S   K   Y   V   E   A   H   Q   Y
  (original)    ATA CCC AGA GAT GAT GGA GTA GAA TGC CCA   GTG ACC TTG CTT TAT CCT TTA TTA TCG GAT   AAA TCA AAA TAC GTC GAG GCT CAC CAA TAT
  (mammalian)   ATC CCC AGA GAT GAC GGC GTG GAG TGC CCA   GTG ACC CTG CTG TAC CCT CTG CTG AGC GAC   AAG AGC AAG TAC GTG GAG GCC CAC CAG TAC
                **  *** *** *** **  **  **  **  *** ***   *** ***  ** **  **  ***  *   *      **    **      **  *** **  *** **  *** **  **

mBaoJin          T   I   C   K   P   L   H   N   Q   P       A   P   D   V   P   Y   H   W   I   R       K   Q   Y   T   Q   S   K   D   D   A
  (original)    ACA ATC TGC AAG CCT CTT CAT AAT CAA CCA   GCA CCT GAT GTC CCA TAT CAC TGG ATT CGT   AAA CAA TAC ACA CAA AGC AAA GAT GAT GCC
  (mammalian)   ACC ATC TGC AAG CCC CTG CAC AAT CAG CCA   GCC CCC GAT GTG CCA TAC CAC TGG ATC AGA   AAG CAG TAC ACC CAG AGC AAG GAC GAC GCC
                **  *** *** *** **  **  **  *** **  ***   **  **  *** **  *** **  *** *** **   *    **  **  *** **  **  *** **  **  **  ***

mBaoJin          E   E   R   D   H   I   C   Q   S   E       T   L   E   A   H   L   K   G   M   D       E   L   Y   K   *
  (original)    GAG GAA CGC GAT CAT ATC TGT CAA TCA GAG   ACT CTC GAA GCA CAC TTA AAG GGC ATG GAC   GAG CTG TAT AAG TAG
  (mammalian)   GAG GAG AGA GAC CAC ATC TGC CAG AGC GAG   ACC CTG GAG GCC CAC CTG AAG GGC ATG GAC   GAG CTG TAC AAG TAA
                *** **   *  **  **  *** **  **      ***   **  **  **  **  ***  *  *** *** *** ***   *** *** **  *** **
```

**Supplementary Fig. 2**

*top,* Nucleotide sequence alignment of mStayGold (E138D) having bacterial-preferred codons and mStayGold(B) having mammalian-preferred codons.

*bottom*, Nucleotide sequence alignment of mBaoJin having jellyfish-original codons and mBaoJin having mammalian-preferred codons.

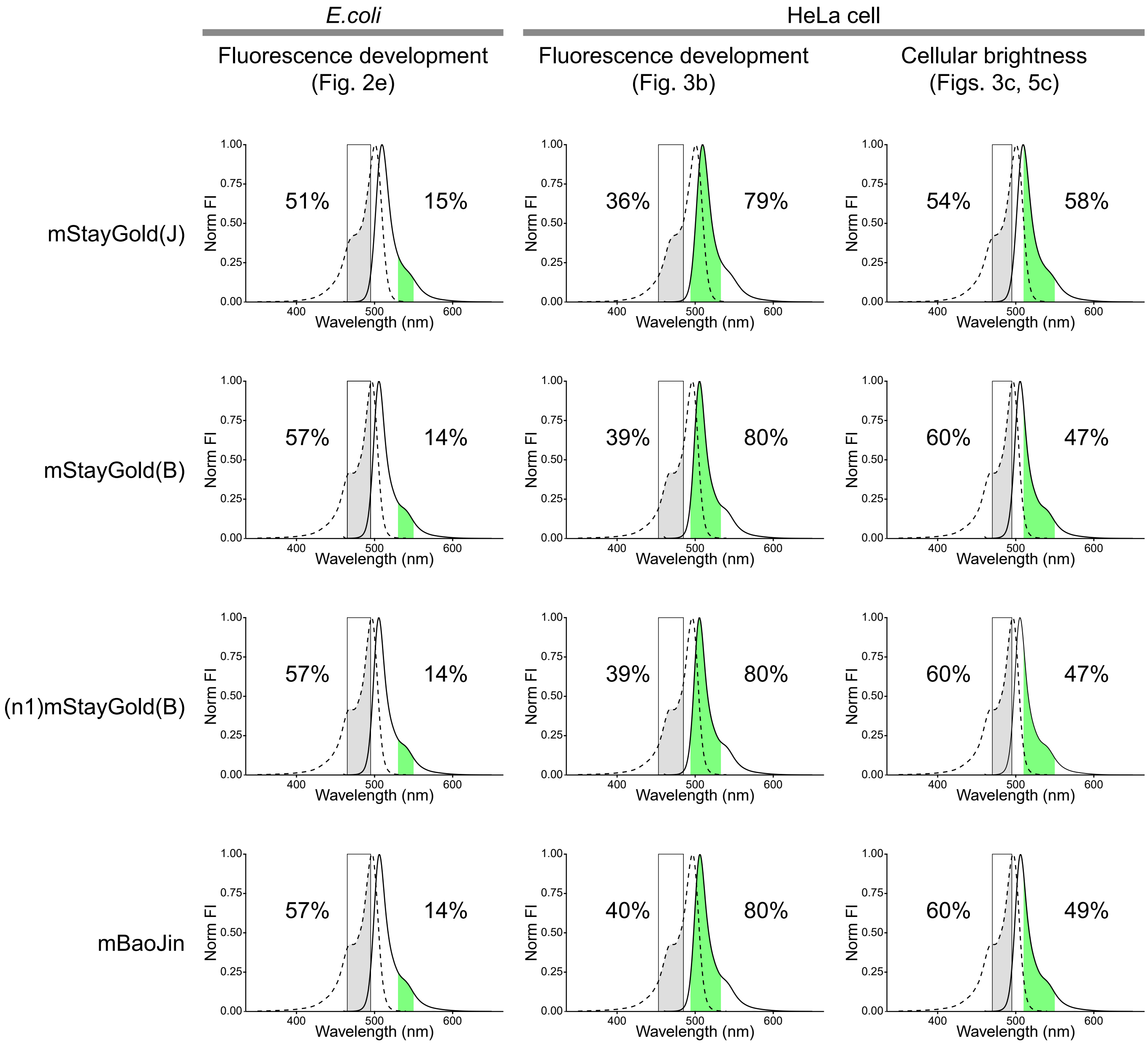


**Supplementary Fig. 3**

Spectral throughputs of StayGold monomers in the imaging systems for observation of fluorescence development (Figs. 2e and 3b) and for determination of cellular brightness (Figs. 3c and 5c). Normalized excitation (dotted line) and emission (solid line) spectra of individual mSGs and transmissions occurring in the excitation (gray) and emission (light-green) passbands. Relative excitation and emission detection efficiencies are shown on the left and right sides of the spectra, respectively. Their products were used to calculate the correction factors. FI: fluorescence intensity.

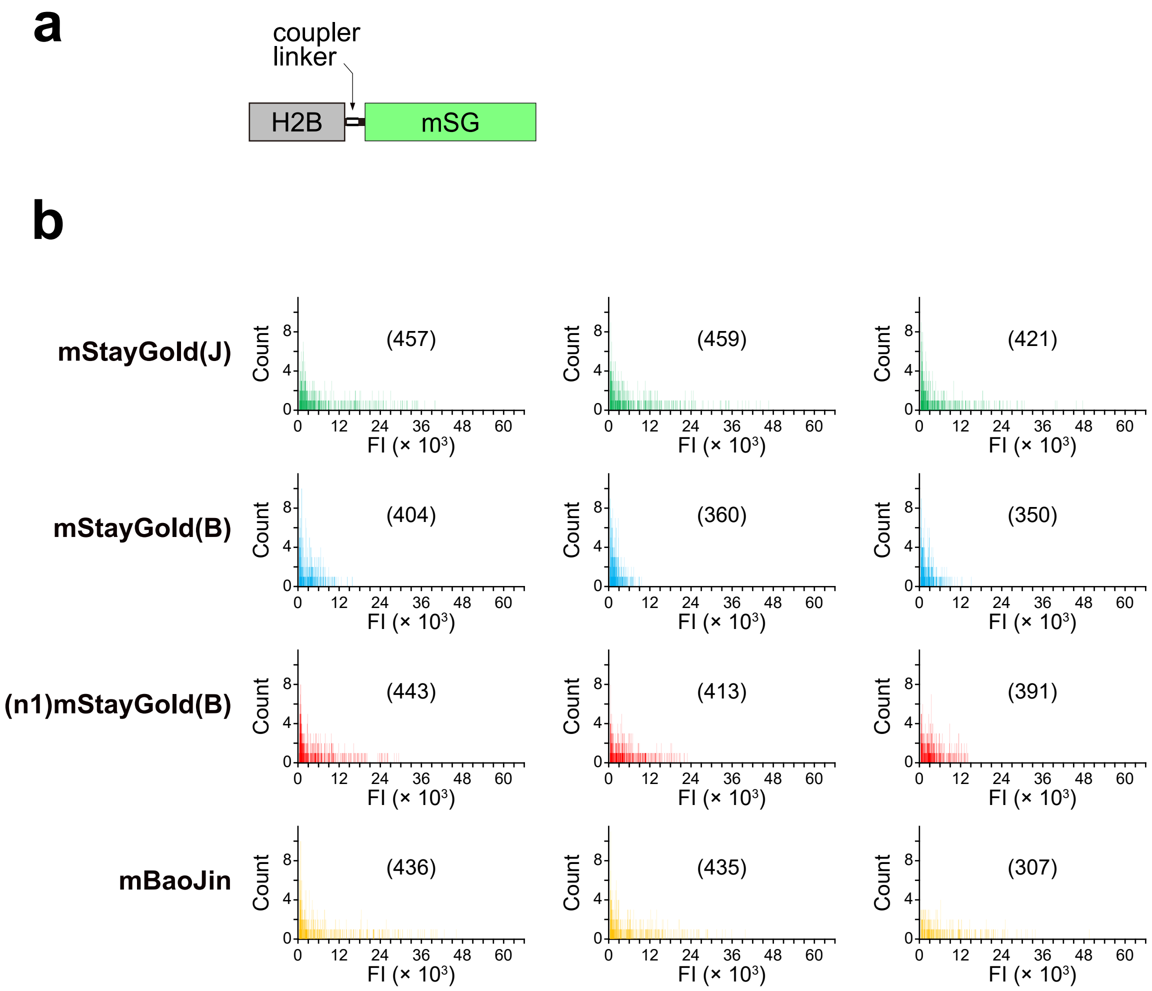


**Supplementary Fig. 4**

Brightness of StayGold monomers in the nucleus of live HeLa cells.

**a,** Domain structure of H2B-FP. H2B: histone 2B. mSG: StayGold monomer.

**b,** Fluorescence intensity (FI) histograms. Sixteen-bit [0, 65,535] intensity images were analyzed. Side-by-side transfection was repeated three times. The number of analyzed cells is shown in the parentheses.

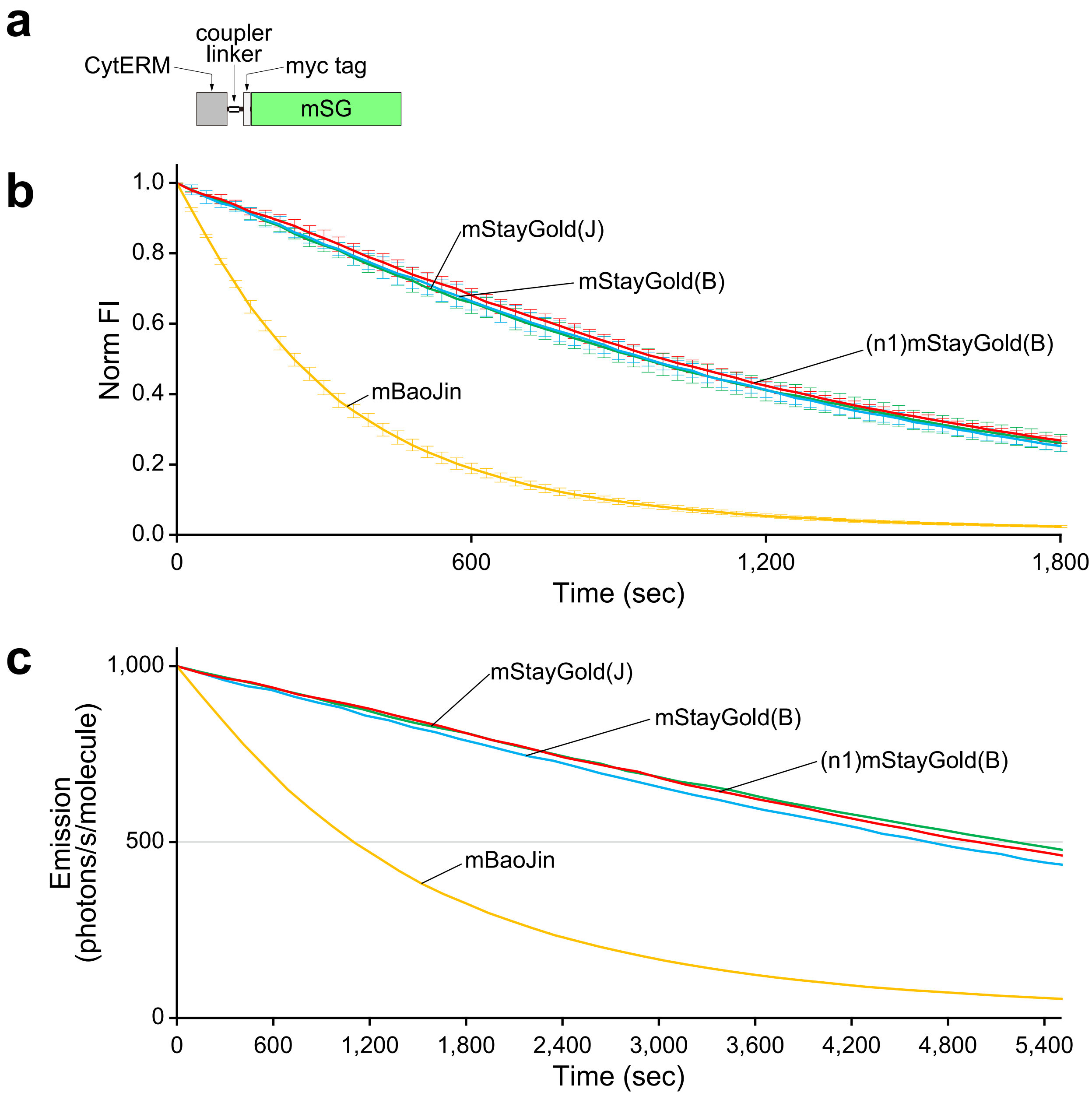


**Supplementary Fig. 5**

Photostability of StayGold monomers anchored to the ER membrane in live HeLa cells.

**a,** Domain structure of CytERM=myc-mSG. "=" denotes "Coupler linker," a triple repeat of the amino acid linker: Gly-Gly-Gly-Gly-Ser (ref. 3). mSG: StayGold monomer.

**b,** Intensity-normalized curves. FI: fluorescence intensity.

**c,** Plot of intensity vs. normalized total exposure time, with an initial emission rate of 1,000 photons $s^{-1}$ $molecule^{-1}$. Illumination intensity, 8.66 W $cm^{-2}$.

**b, c,** The curves are representative of three repetitions ($n$ = 3 independent experiments).

Error bars indicate s.d. (**b**). The statistical values of $t_{1/2}$ (time for photobleaching from an initial emission rate of 1,000 photons $s^{-1}$ $molecule^{-1}$ down to 500 photons $s^{-1}$ $molecule^{-1}$) are shown in Table 1. Traces are indicated in green for mStayGold(J), aqua for mStayGold(B), red for (n1)mStayGold, and orange for mBaoJin.

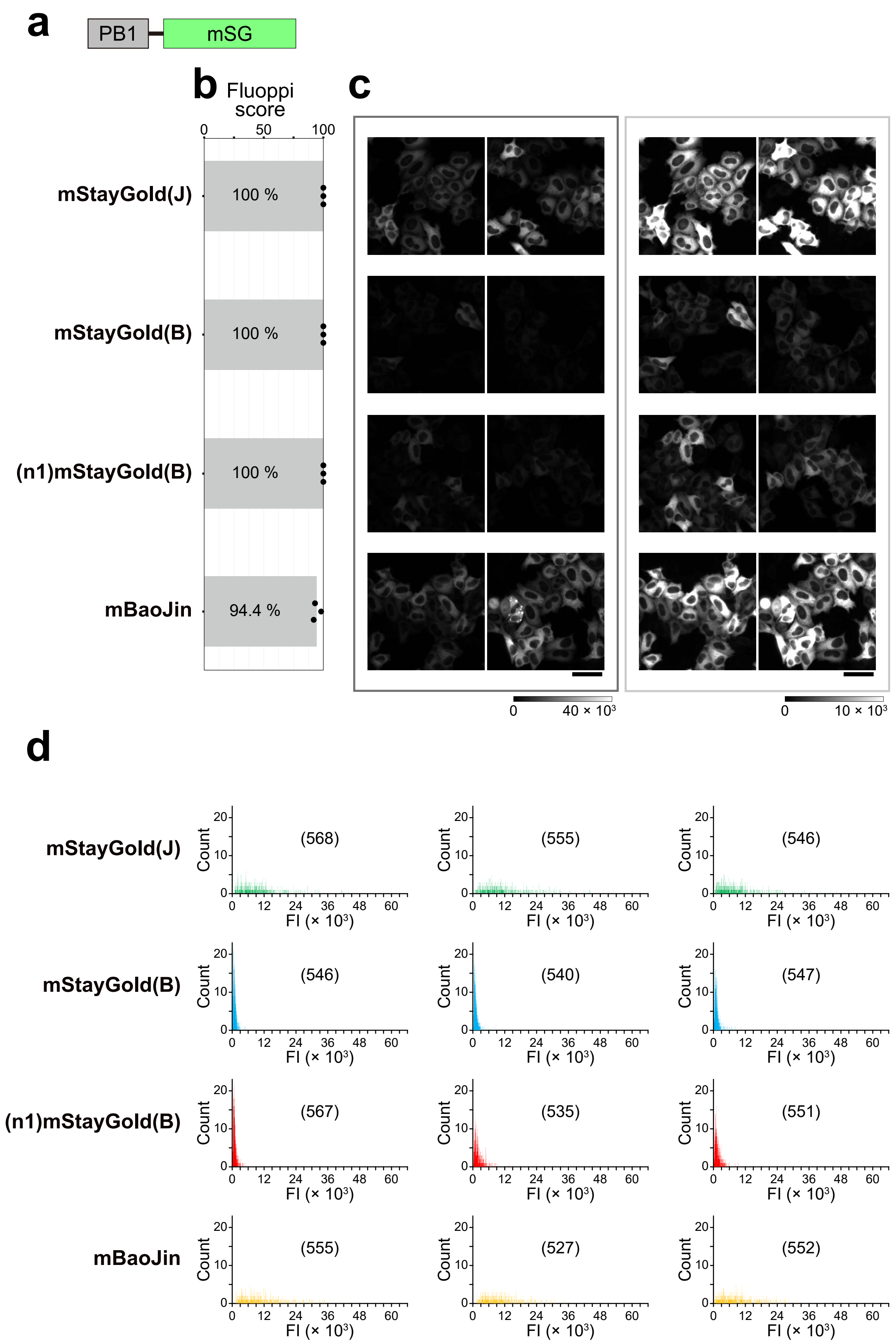


**Supplementary Fig. 6**

Fluoppi assay for assessment of dispersibility and brightness of StayGold monomers.

**a,** Domain structure of PB1-mSG. mSG: StayGold monomer.

**b,** Dispersibility. The percentage of cells scored without fluorescent puncta is plotted (black dots).

**c,** Two representative close-up images are shown with different ranges of display brightness for each construct; Gray scales indicate the lowest and highest intensities of the fluorescence images.

**d,** Brightness in the cytoplasmic compartment. Fluorescence intensity (FI) histograms. Sixteen-bit [0, 65,535] intensity images were analyzed. Side-by-side transfection was repeated three times. The number of analyzed cells is shown in the parentheses.

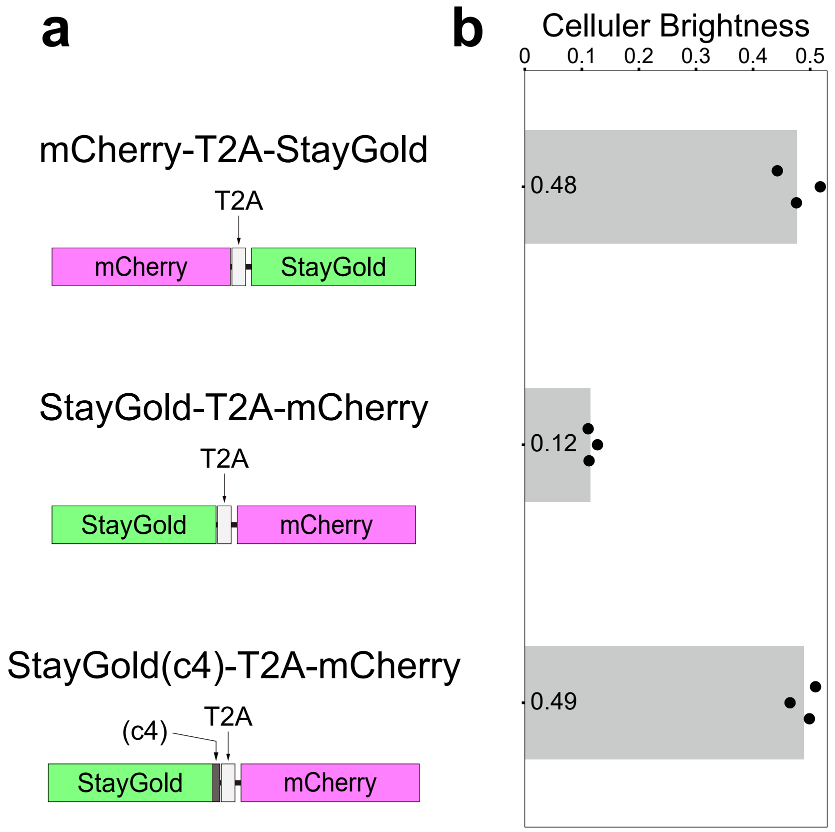


**Supplementary Fig. 7**

Brightness of StayGold in live HeLa cells.

**a,** Cotranslation of StayGold with mCherry using the bicistronic coexpression system. Transfection was performed with pCSII-EF/mCherry-T2A-StayGold (top), pCSII-EF/StayGold-T2A-mCherry (middle) or pCSII-EF/StayGold(c4)-T2A-mCherrry.

**b,** Cellular brightness 48 h after transfection. The green fluorescence was corrected for the mCherry fluorescence. Transfection was repeated three times; mean values are shown by gray bars with plotted dots.

In our previous study[1,2], we determined the cellular brightness of StayGold by using a construct: mCherry-T2A-StayGold. We recently noticed that another cotranslation system in the reversed order was used for this purpose. One example is seen in the study in which mBaoJin was developed[4]. Another example is pDRF-GW(n1)StayGold-T2A-mCherry (addgene). In these cases, however, StayGold must be followed by a c4 adaptor to generate StayGold(c4)-T2A-mCherry. Otherwise, StayGold cannot be expressed well and its cellular brightness will be considerably underestimated. In fact, we confirmed that the high cellular brightness was obtained by StayGold(c4)-T2A-mCherry as well as mCherry-T2A-StayGold but not by StayGold-T2A-mCherry.

pcDNA3/F-tractin=mStayGold and pcDNA3/mStayGold(c4)=UtrCH (ref. 1) are now available from addgene or RIKEN BRC. They can be used for making POI=mStayGold and mStayGold(c4)=POI, respectively, by simple gene exchange. “=” denotes “Coupler linker” (ref. 3).
